# PlotGDP: an AI Agent for Bioinformatics Plotting

**DOI:** 10.64898/2026.01.31.702995

**Authors:** Xiaotong Luo, Ying Shi, Hao Huang, Hongge Wang, Wenyi Cao, Zhixiang Zuo, Qi Zhao, Yueyuan Zheng, Yubin Xie, Shuai Jiang, Jian Ren

**Author notes:** Correspondence to: Xiaotong Luo; Shuai Jiang; Jian Ren. Xiaotong Luo, Ying Shi, Hao Huang, and Hongge Wang contributed equally to this work, and wish to be regarded as joint first authors.

## Abstract

High-quality bioinformatics plotting is important for biology research, especially when preparing for publications. However, the long learning curve and complex coding environment configuration often appear as inevitable costs towards the creation of publication-ready plots. Here, we present PlotGDP (https://plotgdp.biogdp.com/), an AI agent-based web server for bioinformatics plotting. Built on large language models (LLMs), the intelligent plotting agent is designed to accommodate various types of bioinformatics plots, while offering easy usage with simple natural language commands from users. No coding experience or environment deployment is required, since all the user-uploaded data is processed by LLM-generated codes on our remote high-performance server. Additionally, all plotting sessions are based on curated template scripts to minimize the risk of hallucinations from the LLM. Aided by PlotGDP, we hope to contribute to the global biology research community by constructing an online platform for fast and high-quality bioinformatics visualization.

Graphical abstract.
PlotGDP: an AI Agent for Bioinformatics Plotting

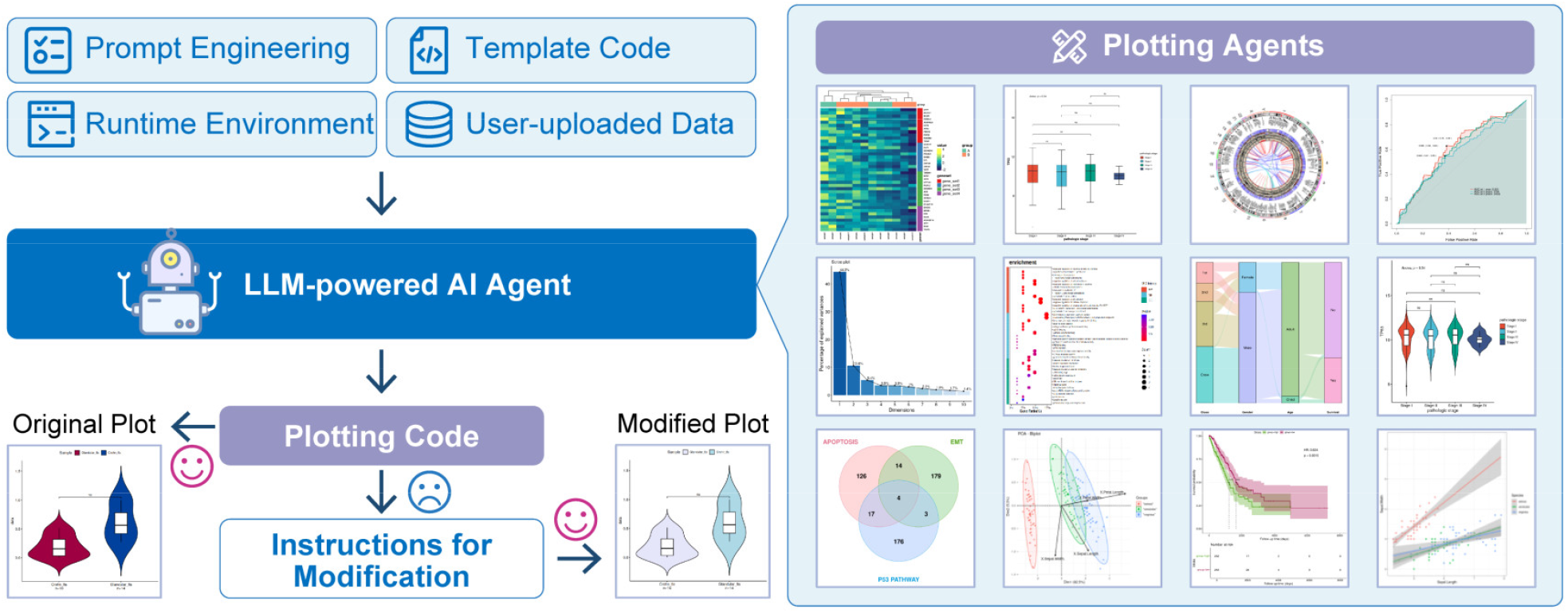

## INTRODUCTION

High-quality bioinformatics plots are essential in biomedical research, offering intuitive and concise representations that enhance the communication of complex findings (1). With the rapid growth of high-throughput sequencing technologies, biomedical data have expanded dramatically, placing bioinformatics analysis at the core of data-driven research (2). For many researchers, generating high-quality figures from data analysis results remains a significant challenge (3). This is particularly true for those without programming experience, who often struggle to apply advanced and widely adopted bioinformatics visualization methods to present their findings effectively (4). Learning to code from scratch for figure generation is both time-consuming and labor-intensive, highlighting the need for more efficient and standardized approaches to biological data visualization.

To address this, several platforms such as GEO2R (5), GEPIA (6), HemI 2.0 (7), and MetaScape (8) have been developed. However, they are typically restricted to a narrow range of plots and cannot fully meet the needs of multi-omics research. In recent years, additional integrated plotting tools have been introduced. For instance, SRplot (9) offers approximately 20 common plotting functions; however, its commercial pricing model is prohibitively expensive for many academic users and research laboratories. Tools like SangerBox 2.0 (10), HiOmics (11), and BioLadder 2.0 (12) provide broader visualization options, yet still require payment for more complex plots. Similarly, tools like Chiplot (https://www.chiplot.online/) offer free plotting functions, but access to certain advanced features or course content remains fee-based. Although these platforms expand visualization capabilities, they often depend on complex preprocessing steps, involve cumbersome parameter settings, and provide limited interactivity, thereby reducing flexibility and efficiency while preventing users from refining details beyond predefined parameters.

Our team has previously developed several widely adopted visualization tools, including Domain Graph (DOG) (13) for protein domain mapping, Illustrator for Biological Sequences (IBS/IBS 2.0) (14,15), and BioGDP (16), a platform for generating biomedical schematic diagrams. These tools allow researchers to create and edit high-quality graphics through simple, user-friendly interactions, and their widespread adoption highlights the value of achieving high-quality biomedical visualizations via streamlined, accessible interfaces. However, they are primarily designed for biomedical illustrations and cannot generate bioinformatics plots directly from data. Therefore, there is an urgent need for a tool that can flexibly produce accurate statistical analysis plots through straightforward interactions.

Recent advances in large language models (LLMs) have created new opportunities for bioinformatics (17). Emerging LLM-based intelligent agents, such as Biomni (18), show potential for enhancing reproducibility and analytical efficiency by automating complex bioinformatics workflows, integrating heterogeneous data sources, and reducing human-induced variability in data processing. However, existing agents remain limited in stability and adaptability. They are primarily designed for complex data analysis tasks and often struggle to handle errors or unexpected situations, while also lacking the ability to adjust graphical outputs flexibly. This underscores the need to develop more powerful and flexible LLM-based approaches for bioinformatic data visualization.

In this study, we developed PlotGDP (https://plotgdp.biogdp.com/), an LLM-powered platform for bioinformatics plotting (**Figure 1**). PlotGDP is a web-based integrated development environment that requires no local deployment. It supports conversational programming, allowing users to automatically generate code through natural language interactions and quickly produce diverse, publication-ready plots with uploaded data. Beyond automated generation, PlotGDP provides a library of diverse, commonly used bioinformatics visualization modules for plotting, learning, reusing, and sharing. Its interactive interface enables real-time adjustments to plots, code, and parameters through either dialogue or manual editing, greatly enhancing usability and flexibility, and helping users efficiently achieve data visualization and presentation. Together with its sister product BioGDP, PlotGDP forms a complementary ecosystem that addresses key visualization needs in bioinformatics, spanning data-driven statistical plots and schematic biomedical illustrations.

**Figure 1.**
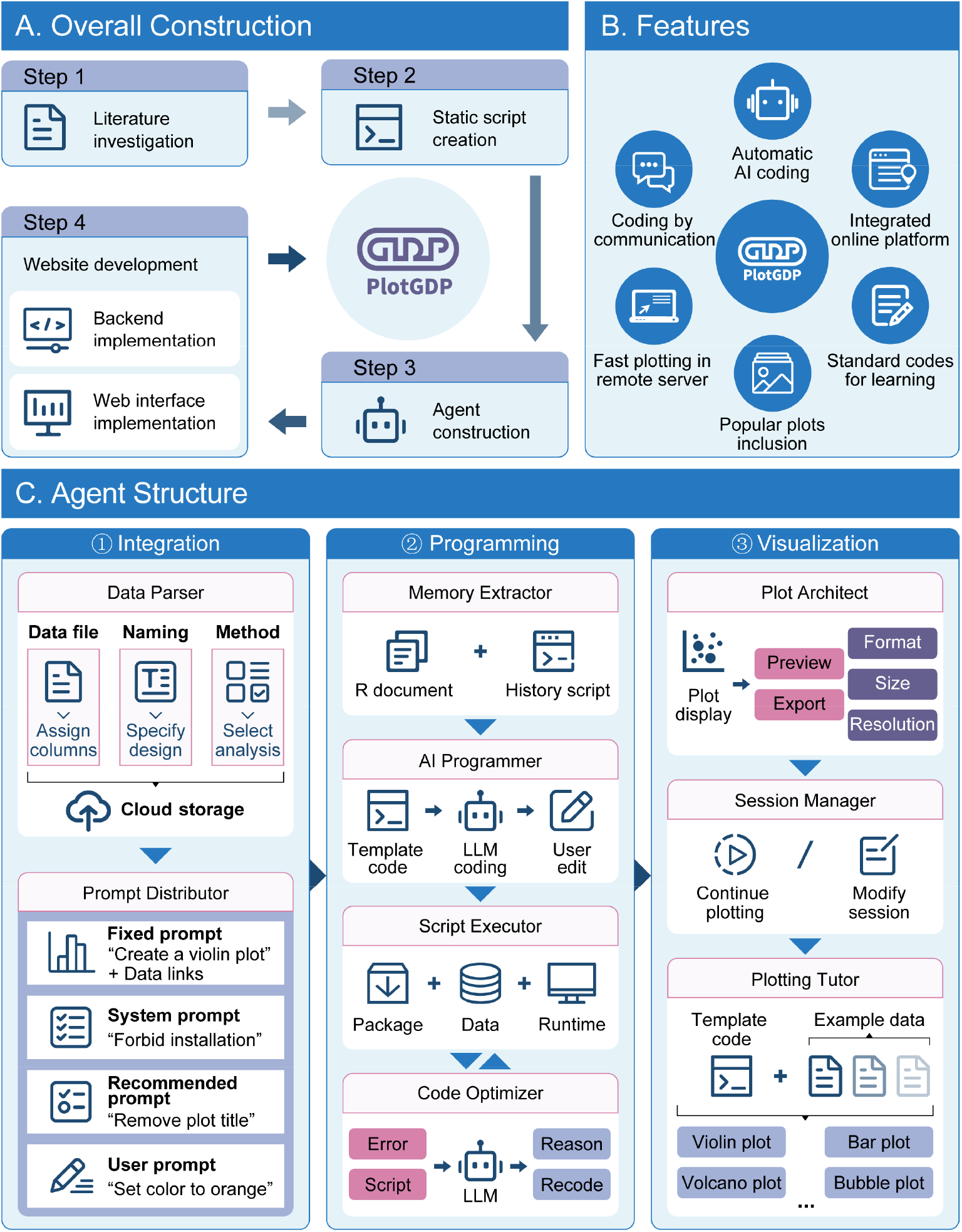
Overview of PlotGDP. **(A)** Overall construction of PlotGDP. 4 steps were involved in the construction. **(B)** Features of PlotGDP. 6 key features are covered. **(C)** Agent structure of PlotGDP. 3 sections are integrated.

## MATERIALS AND METHODS

The overall implementation of PlotGDP is presented in **Figure 1A**.

### Plotting Code Organization and Compilation

To start with, various plots that are frequently used for visualization in biology research according to the literature were determined: heatmap, Venn diagram, Sankey plot, boxplot, violin plot, scatter plot, Kaplan-Meier (KM) survival curve, survival ROC plot, differentially expressed gene (DEG) volcano plot, circos plot, principal component analysis (PCA) scree plot, PCA biplot, and GO/KEGG enrichment bubble plot/bar plot. Widely accepted R packages for plotting were integrated into the corresponding template code, including ggplot2 (v3.5.2) (19), pheatmap (v1.0.13), VennDiagram (v1.7.3) (20), ggalluvial (v0.12.5) (21), ggpubr (v0.6.0), survminer (v0.5.0), survival (v3.8-3) (22), survivalROC (v1.0.3.1), RCircos (v1.2.1) (23), and factoextra (v1.0.7). For each type of plot, we compiled a specific template R script as a coding reference for the LLM, in order to minimize the risk of coding hallucination. These template scripts were displayed on the “Code Hub” of PlotGDP. Specifically, the R script of each plot type containing template codes was encapsulated into a standardized function for users to invoke. These functions integrated diverse plotting-associated analytical workflows for the processing of raw data, besides the plotting codes. All the packages and codes were compiled and fully tested on our server with the same environment for future user access.

### LLM prompting

To enable precise bioinformatics plotting through code modification, we developed a comprehensive LLM prompting strategy consisting of three components, besides user-entered prompts.

a. Fixed prompts: serve as the initial instructions at the start of each plotting session, guiding the LLM according to the plot type selected by the user, for example, “please provide the code for plotting a violin plot”. Data links are also integrated.
b. System prompts: provide explicit instructions for the LLM to generate user-customized plotting codes based on the template scripts, ensuring executability while avoiding hallucinations and preventing improper actions, such as installing inappropriate packages.
c. Recommended prompts: offer example commands that allow users to request modifications of existing code, enabling flexible adjustment of the plotting details, such as “change the legend title to ‘PCA legend’”. These examples demonstrate practical prompting patterns for users to learn from and adapt.

All prompts, together with user-uploaded data, user-defined parameters, and template code, were processed through the Qwen3 (qwen3-coder-plus) LLM application programming interface (API) provided by Alibaba Cloud (https://www.aliyun.com/product/tongyi).

### AI agent construction

We designed and implemented a comprehensive AI agent to interact with the LLM for automated plot generation. At the beginning of each plotting session, user-defined parameters are integrated with system prompts, fixed instruction prompts, and template code to construct a unified input prompt. This prompt is then submitted to the LLM together with official R documentation from relevant visualization packages as contextual references. The input parameters include strings for plotting variable names, method selections, and directory paths associated with user-uploaded data. The uploaded tabular files are required to contain columns that conform to the data extraction logic specified in the template code. Executable example datasets are also provided for user reference and validation.

Upon receiving the response of the LLM, the agent programmatically extracts both the generated R code and the accompanying natural language explanations. The code is executed, and the resulting plots are rendered and presented to the user. During iterative refinement, the agent follows the same workflow while incorporating additional user feedback to update the existing script. Each subsequent output incrementally modifies the code generated in the previous iteration, enabling controlled and traceable script evolution.

This execution step is supported by an integrated R compilation module that provides a controlled and reproducible runtime environment for LLM-generated code. In addition, the AI agent is seamlessly connected with auxiliary system components, including data storage, file upload and download services, and directory management, thereby enabling a complete end-to-end plotting workflow.

### Backend and web interface implementation

We employed a hybrid data storage architecture that combines MySQL and MongoDB. The LLM’s capabilities are invoked via RESTful APIs. The backend coding was conducted in Java and SpringBoot 3.0 microservice framework. The frontend was developed with the React framework, integrating Hypertext Markup Language (HTML), Cascading Style Sheets (CSS), and JavaScript (JS), to deliver a responsive and user-friendly interface.

### Application of PlotGDP in cancer research

To showcase the real-world application, we applied PlotGDP on a Gene Expression Omnibus (GEO) (5) human breast cancer case study with accession “GSE37751”. The gene expression data, survival data, and grouping information were analyzed and visualized in 6 plotting modules from PlotGDP: DEG volcano plot, GO/KEGG enrichment bubble plot, Venn diagram, PCA biplot, KM survival curve, and violin plot. Gene-term associations from PubMed literature were obtained from Geneshot (24).

## RESULTS

### Overview of the advantages of PlotGDP

Six major features distinguish PlotGDP from other bioinformatic plotting tools (**Figure 1B, Table 1**): (a) PlotGDP can automatically generate production-ready code with the LLM, minimizing researchers’ learning costs. (b) Dozens of widely used bioinformatic plot types are integrated to cover key visualization needs. (c) A fully integrated online platform enables convenient plotting, eliminating the need for local environment setup. (d) No prior programming experience is required; users can create and modify publication-quality plots simply by providing brief natural language instructions to the AI agent. (e) Standardized codes serve as an educational resource for beginners to learn bioinformatic plotting scripts, while also feeding the LLM as executable coding references. (f) A regular laptop suffices for plotting functionalities, as all computations are performed on our high-performance server.

**Table 1.** Feature comparison for bioinformatic plotting tools. ^a^.

| Feature | PlotGDP | Biomni | OLAF | HiOmics | Hiplot | ImageGP 2.0 | SangerBox | SRplot | BioLadder 2.0 | Chiplot |
| --- | --- | --- | --- | --- | --- | --- | --- | --- | --- | --- |
| **Major feature** |  |  |  |  |  |  |  |  |  |  |
| AI-driven coding | √ | √ | √ | × | × | × | × | × | × | × |
| Availability of template codes | √ | × | × | × | × | × | × | × | × | × |
| Running online | √ | √ | × | √ | √ | √ | √ | √ | √ | √ |
| Natural language communication | √ | √ | √ | × | × | × | × | × | × | × |
| **Additional feature** |  |  |  |  |  |  |  |  |  |  |
| Debugging | √ | √ | √ | × | × | × | × | × | × | × |
| Free of charge | √ | - | √ | - | - | √ | - | × | - | - |
| Fast response | √ | × | - | √ | √ | √ | √ | √ | √ | √ |
| Direct code editing | √ | × | × | × | × | × | × | × | × | × |
| Explanation for parameters | √ | × | × | √ | √ | √ | √ | √ | √ | √ |
| Example data | √ | × | × | √ | √ | √ | √ | √ | √ | √ |
| AI-driven Q&A | √ | √ | √ | × | × | × | × | × | √ | × |

<sup>a</sup> Symbol explanation: "√" means "yes", "×" means "no", "-" means "depending on the usage".

Additional features (**Table 1**) include continuous debugging, free usage, parameter explanations, and example datasets, distinguishing PlotGDP from similar tools such as Biomni (18), OLAF (25), HiOmics (11), Hiplot (26), ImageGP 2.0 (27), SangerBox (10), SRplot (9), BioLadder 2.0 (12), and Chiplot (https://www.chiplot.online/). Relatively, the solidity and flexibility of PlotGDP make it a more appropriate tool for bioinformatic visualization.

### Agent structure of PlotGDP

PlotGDP contains a comprehensive agentic workflow with 3 sections (**Figure 1C**): (a) Integration. The “data parser” first processes files, naming, and methods specified by users, and stores them on the cloud. Their information contributes to the fixed prompt, along with the system prompt, the recommended prompt, and the user prompt, arranged by the “prompt distributor”. (b) Programming. The “memory extractor” conveys the information of R documents and history scripts to the “AI programmer” for LLM-driven coding. The template codes and the aforementioned prompts are used as coding guidance. After the user edits, the “script executor” runs the code in the remote runtime environment with the given packages and specified data. If errors appear, they will be analyzed by the “code optimizer” with LLM-powered recoding, and rerunning afterwards. (c) Visualization. The “plot architect” displays the plot from the executor, supporting quantitative preview and export. For completed plotting tasks, the “session manager” supports continued plotting or session modification. Finally, the “plotting tutor” demonstrates example plotting sessions with template codes and example data for users to learn.

### Web interface and functionality

The visualization journey in PlotGDP starts from the “plotting agents”, which is a gallery of plotting bots for the most widely used bioinformatic visualizations (**Figure 2A**). In the current implementation, PlotGDP supports 14 commonly used plot types, including heatmaps, volcano plots, violin plots, bubble enrichment plots, bar enrichment plots, circos plots, Sankey plots, survival ROC plots, scatter plots, Venn plots, survival curves, box plots, scree plots, and PCA biplots. Each plotting agent is accompanied by structured plot descriptions and search functionality to facilitate efficient discovery. Upon selecting a plotting agent, users are directed to a new Workspace session in which all parameters and settings are automatically configured to match the selected plot type.

**Figure 2.**
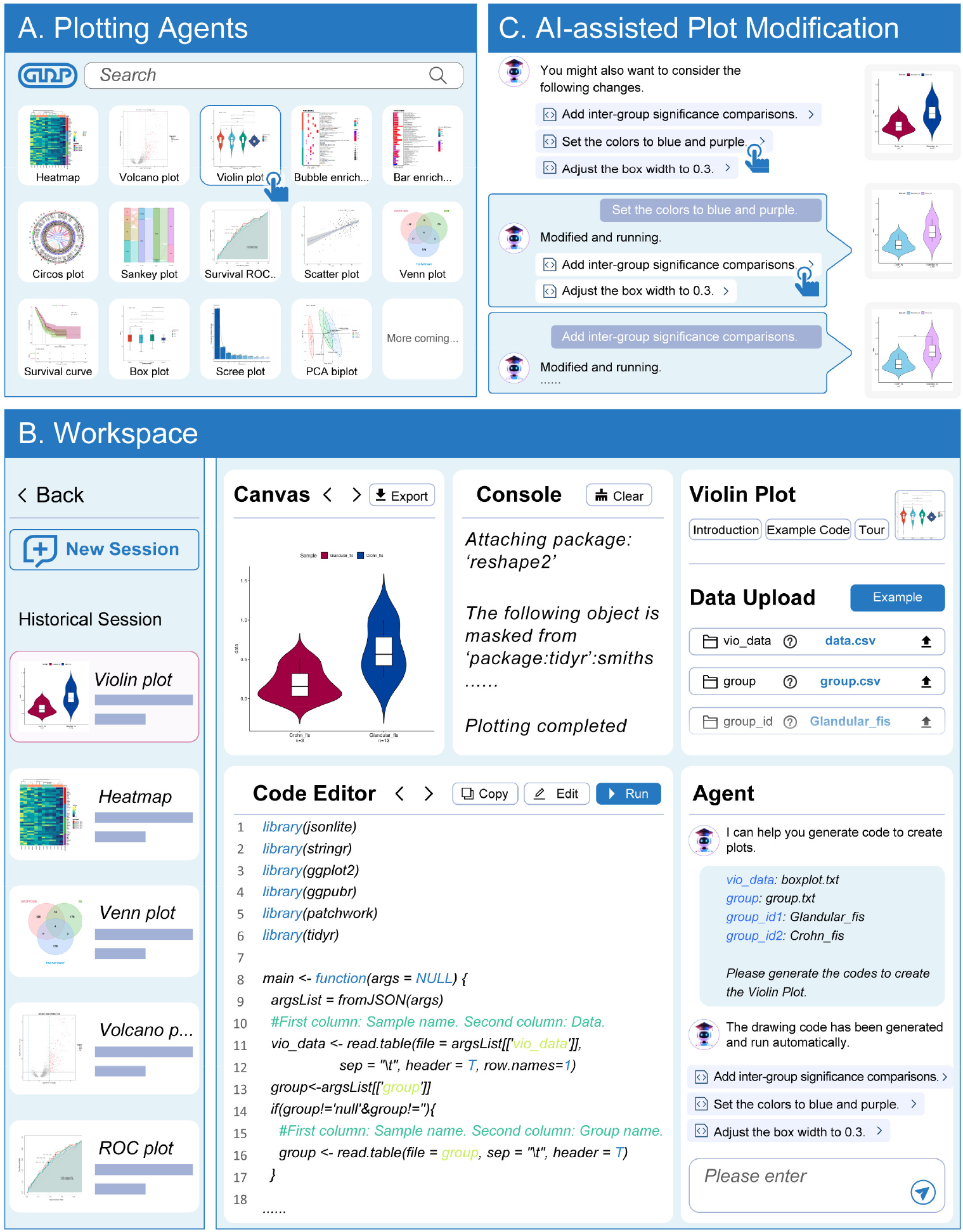
Major web interfaces and functionalities of PlotGDP. **(A)** Plotting agents: the interface to browse, search, and select the intended plotting templates. **(B)** Workspace: the primary webpage of PlotGDP for users to create/modify LLM-generated plots. **(C)** AI-assisted plot modification.

“Workspace” is the primary webpage for users to interact with the plotting agent (**Figure 2B**). At the start of a “New Session” (upper left), users can upload data to the cloud that mimics the structures of example data provided (upper right). These data, together with user-defined parameters and the fixed prompt, are integrated as the input (right bottom) for the LLM to generate the first round of output plotting codes in each session (middle bottom). If precise editing is needed, users can directly modify the LLM-generated code in “Code Editor”. By clicking the “Run” button, the LLM-generated codes will remotely run and return the output plot for users to view the details or download in different formats (upper middle). All sessions are summarized in the sidebar (left), where users can retrieve any history codes from the cloud and continue the edits/plotting. Additionally, detailed introduction and template codes for the chosen plot type are available for users (upper right). In principle, the largest control is given to users to handle plots, files, codes, sessions, etc., with convenient actions like uploading, downloading, removing, renaming, editing, and copying.

As the highlight of PlotGDP, users can describe their needs in natural language as subsequent inputs for the agent to adjust the plot automatically (**Figure 2C**), which guides the LLM to modify the previously generated codes and present new codes to plot. Recommended inputs are displayed for users to click, serving as simple and straightforward prompting examples.

### Case study of PlotGDP

To illustrate the capabilities of PlotGDP in supporting biomedical publications, we present a case study featuring plots generated by PlotGDP (**Figure 3**). These plots provide a concise yet comprehensive overview of the potentially key gene *CCNE2* in breast tumor progression, as detailed below.

**Figure 3.**
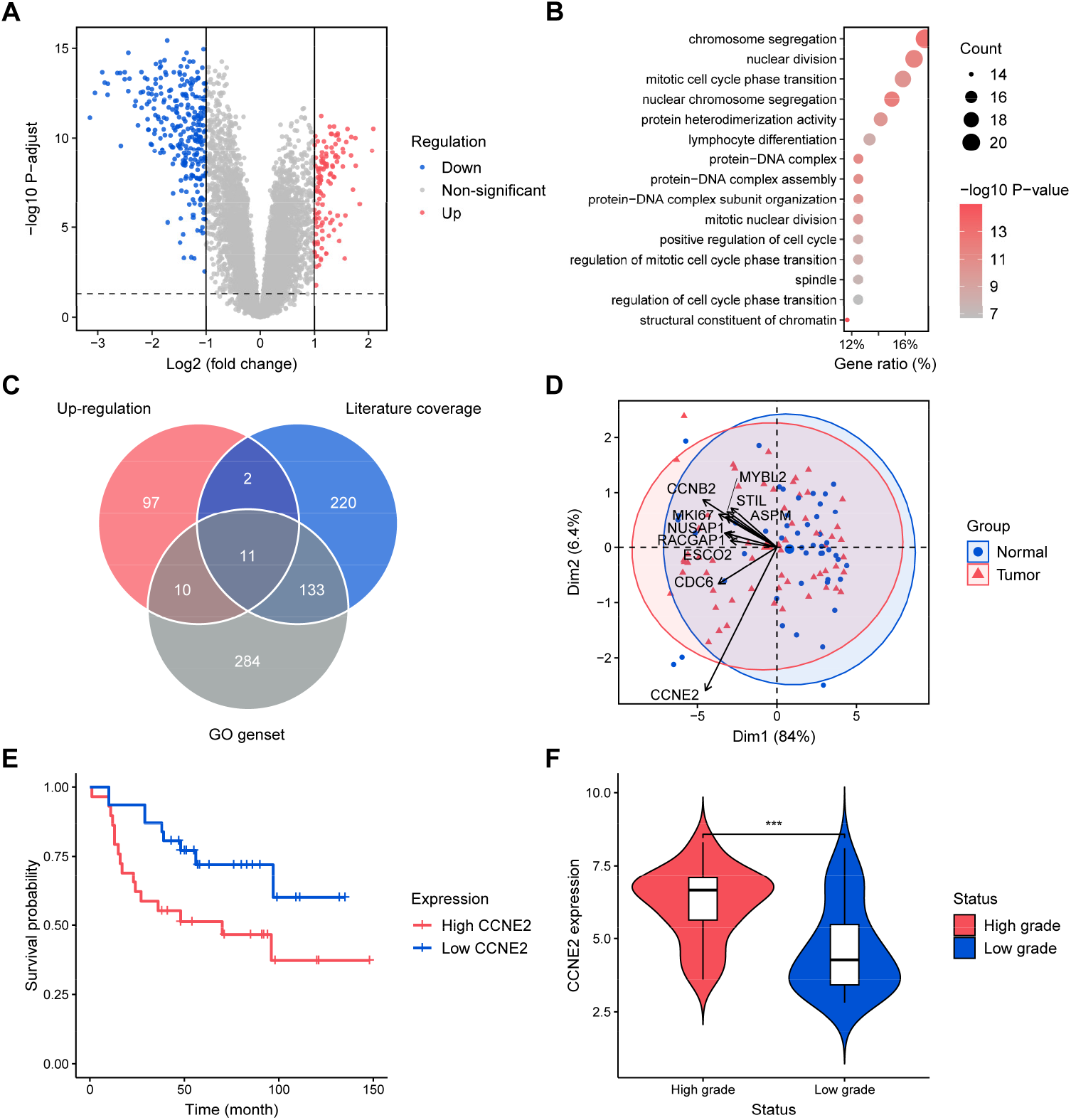
A case study of PlotGDP in breast cancer research. **(A)** A volcano plot of DEGs between the breast tumor and the normal tissue. Significantly up-regulated DEGs are marked in red (log_2_(FC) > 1, P-adjusted < 0.05), while the significantly down-regulated in blue (log_2_(FC) < -1, P-adjusted < 0.05). **(B)** A bubble plot of the most enriched GO terms according to the gene ratio induced by the significantly up-regulated DEGs in the breast tumor. **(C)** A Venn diagram for the numbers of genes with 3 components: significantly up-regulated genes in the breast tumor (red), genes related to “chromosome segregation” in previous PubMed literature according to Geneshot (blue), and genes involved in “chromosome segregation” GO geneset (grey). **(D)** A PCA plot for the breast tumor and normal tissue. The tumor samples are in red, while the normal samples are in blue. **(E)** A survival plot for breast cancer patients with high *CCNE2* expression (red) and low *CCNE2* expression (blue). **(F)** A violin plot for the expression of *CCNE2* in high-grade breast tumors (red) and low-grade breast tumors (blue). ***: p ≤ 0.001, Wilcoxon test for the mean comparison of p-values.

Starting with the differential analysis of a transcriptomic dataset (GSE37751 from GEO), PlotGDP identified 400 DEGs, in which 120 were significantly up-regulated in breast tumor samples compared to normal tissues (**Figure 3A**). The up-regulated DEGs were enriched in several GO terms, the strongest of which was “chromosome segregation” according to PlotGDP (**Figure 3B**). By searching for this GO term in Geneshot (24) to obtain a list of relevant functional genes that were already studied in existing PubMed literature, we identified 10 unstudied genes that are both significantly up-regulated in breast tumors and relevant to chromosome segregation, based on the “Venn diagram” from PlotGDP (**Figure 3C**). *CCNE2*, among these 10 genes, contributes the most to the variance according to the PlotGDP-derived “PCA biplot” (**Figure 3D**), which is then selected as a potential tumor-driver gene for further PlotGDP visualizations. The survival curve for patients with higher expression of *CCNE2* shows a faster decline in the survival probability compared to those with lower expression (**Figure 3E**), and the expression of *CCNE2* seems to increase with the progression of the breast tumor (**Figure 3F**) as well. Both pieces of stratified evidence from PlotGDP revealed the potential contribution of *CCNE2* to the adverse outcomes in breast cancer patients that call for deeper low-throughput investigation.

## DISCUSSION

In recent years, biomedical research has seen rapid growth in both publication volume and concerns over quality (28), highlighting the need for advanced methods to generate publication-ready plots quickly, conveniently, and accurately. To address this, we developed PlotGDP, an LLM-powered AI agent for efficient bioinformatics plotting across diverse types of commonly used visualizations. PlotGDP enables automatic code generation and modification through a combination of built-in and user-provided prompts, while also offering standardized plotting scripts for learning and a user-friendly interface optimized for backend agents. Users can upload data, specify visualization needs in natural language, remotely execute LLM-generated code, and download high-quality plots: all within seconds.

The key advantage of PlotGDP over other bioinformatics plotting tools lies in its ability to directly translate natural language instructions into executable plotting code, ensuring both accuracy and flexibility. This design eliminates the need for coding skills, allowing researchers to focus on scientific questions rather than technical barriers in data visualization. To further streamline the process, PlotGDP integrates template codes to guide the LLM, leverages fast remote servers, and remains entirely free of charge for all use cases (**Table 1**). However, despite efforts to restrict LLM behaviors, some generated code may still contain errors due to user requests that cannot be fully met (e.g., mismatched column names, missing network connections, or outdated packages). Future versions of PlotGDP will address these by incorporating more precise LLM-driven error explanations and code version management.

To demonstrate the practical utility of PlotGDP in research, we applied it to a cancer key gene discovery study and generated a series of publication-ready plots. (**Figure 3**). These plots followed a strict logic, showcasing a clear workflow, demonstrating target gene discovery from high-throughput transcriptomic data. Beyond oncology, PlotGDP has broad applicability across biomedical research, including cellular biology, tissue physiology, and epidemiology. With ongoing upgrades, PlotGDP aims to expand coverage of plotting types while refining the LLM response pipeline to meet the diverse visualization needs of all biological subfields. Ultimately, our long-term goal is to provide the biomedical research community with a comprehensive plotting platform that enables the generation of high-quality, reproducible visualizations.

## DATA AVAILABILITY

PlotGDP is publicly accessible at https://plotgdp.biogdp.com/. This website is free and open to all users and there is no login requirement.

## FUNDING

This study was supported by the National Natural Science Foundation of China [32470709, 82573769, 32570765, 32570794, 82573731].

## CONFLICT OF INTEREST

None declared.

## ACKNOWLEDGEMENTS

*Authors contributions*: **Xiaotong Luo**: Funding acquisition, Project administration, Writing— review & editing. **Ying Shi**: Methodology, Software, Writing—review & editing. **Hao Huang**: Methodology, Software, Validation. **Hongge Wang**: Methodology, Visualization, Writing— original draft. **Wenyi Cao**: Methodology, Software, Data curation. **Zhixiang Zuo**: Funding acquisition, Project administration, Validation. **Qi Zhao**: Funding acquisition, Project administration, Validation. **Yueyuan Zheng**: Funding acquisition, Project administration, Validation. **Yubin Xie**: Funding acquisition, Project administration, Validation. **Shuai Jiang**: Funding acquisition, Project administration, Resources. **Jian Ren**: Conceptualization, Funding acquisition, Project administration, Resources.

## Notes

https://plotgdp.biogdp.com/

### Competing Interest Statement

The authors have declared no competing interest.

### Summary of Updates

Updated a hyperlink in the abstract.

